# Visual Perception of 3D Space and Shape in Time - Part II: 3D Space Perception with Holographic Depth

**DOI:** 10.1101/2022.02.28.482181

**Authors:** Isabella Bustanoby, Andrew Krupien, Umaima Afifa, Benjamin Asdell, Michaela Bacani, James Boudreau, Javier Carmona, Pranav Chandrashekar, Mark Diamond, Diego Espino, Arnav Gangal, Chandan Kittur, Yaochi Li, Tanvir Mann, Christian Matamoros, Trevor McCarthy, Elizabeth Mills, Stephen Nazareth, Justin Nguyen, Kenya Ochoa, Sophie Robbins, Despoina Sparakis, Brian Ta, Kian Trengove, Tyler Xu, Natsuko Yamaguchi, Christine Yang, Eden Zafran, Aaron P. Blaisdell, Katsushi Arisaka

## Abstract

Visual perception plays a critical role in navigating 3D space and extracting semantic information crucial to survival. Even though visual stimulation on the retina is fundamentally 2D, we seem to perceive the world around us in vivid 3D effortlessly. This reconstructed 3D space is allocentric and faithfully represents the external 3D world. How can we recreate stable 3D visual space so promptly and reliably?

To solve this mystery, we have developed new concepts **MePMoS** (Memory-Prediction-Motion-Sensing) and **NHT** (Neural Holography Tomography). These models state that visual signal processing must be primarily top-down, starting from memory and prediction. Our brains predict and construct the expected 3D space holographically using traveling alpha brainwaves. Thus, 3D space is represented by the three time signals in three directions.

To test this hypothesis, we designed reaction time (RT) experiments to observe predicted space-to-time conversion, especially as a function of distance. We placed LED strips on a horizontal plane to cover distances from close up to 2.5 m or 5 m, either using a 1D or a 2D lattice. Participants were instructed to promptly report observed LED patterns at various distances. As expected, stimulation at the fixation cue location always gave the fastest RT. Additional RT delays were proportional to the distance from the cue. Furthermore, both covert attention (without eye movements) and overt attention (with eye movements) created the same RT delays, and both binocular and monocular views resulted in the same RTs. These findings strongly support our predictions, in which the observed RT-depth dependence is indicative of the spatiotemporal conversion required for constructing allocentric 3D space. After all, we perceive and measure 3D space by time as Einstein postulated a century ago.

## 1 Introduction

Depth perception is an essential aspect of human vision and is used to determine the distances between objects in an observer’s surroundings and the observer. This becomes an important evolutionary tool in evaluating the location of a predator or food source.

Within the current understanding of depth perception, there are various speculated external and cognitive explanations for how vivid 3D vision is possible. The three-dimensional impact of viewing space through various physical means could include the appearance of motion parallax, occlusion, linear perspective, the elevation effect, and the kinetic depth effect from the moving of objects within viewed space. Various internal factors possibly contributing to the perception of depth are the difference between monocular and binocular viewing, the adjustment of the focal length of the eye’s lens, the triangulation of 3D space from binocular/stereo view and binocular disparity. All of these factors contribute to our vision but understanding how these various strategies possibly form a conclusive comprehension of 3D space from two-dimensional retinal information is unknown.

Binocular disparity and visual triangulation are all mechanisms that rely on binocular vision or the viewer to experience 3D space through their own movement. Binocular disparity refers to the slightly different, offset images captured by both eyes. Stereopsis is the process by which our brain can extract 3D information from binocular disparity. It first needs to determine the absolute disparity, or the difference in angular separation between the two eyes and their foveas, of corresponding points in the images captured by each eye. This process is considered a way that the brain can produce depth from 2D information. Motion parallax uses similar geometry to binocular disparity, as this also gathers information about depth as the observer moves throughout their surroundings while collecting images from different viewpoints. Optical flow refers to the change in light between different visual frames as an observer moves throughout their environment. The rays corresponding to different points and the observer are projected in a regulated and predictable manner. These rays provide information about their angular velocities and are projected onto the retina (Simpson, 1993).

Different neural substrates guide object identification and object location (and also movement). Based on the Two-Stream Hypothesis (Goodale, Milner 1992), there exist the Dorsal and Ventral Pathways. The dorsal pathway, starting from the striate cortex/primary visual cortex to the posterior parietal region, is relevant to this study. It processes information about the spatial location of an object to an observer and provides information about the orientation and shape of objects (Hebart 2012). There’s also an established understanding of the specific regions that make up the visual area of the brain: V1, V2, V3, V4, as established by the Brainnetome Atlas (Fan et al. 2016).

As past research investigates the structure and functionality of the brain, understanding where 3D visual processing is specifically facilitated has been further studied. It was found that recorded neurons in the MT were active while macaque monkeys reacted to visual stimuli, in order to study neural substrates for depth perception (Nadler 2008). It was further observed that the MT plays a role in visual motion perception, as the MT contains neurons that are sensitive to binocular disparity, the area used for stereopsis (Kim 2016).

This previous research is still not enough to understand depth, for flashing neurons at the MT cannot reasonably be responsible for depth alone. As established previously, different depths create different neuronal spike rates, but in the brain the different spike rate does not encode to create the perception of depth. An explanation for this could involve the role of spike timing. Timing matters, if a cell is firing right before another, the cells may be linked together (Markram 2012). Furthermore, synapses increase in strength if presynaptic spikes happen milliseconds before the postsynaptic spike.

Top-Down Processing can be described as higher-order representations that influence earlier information processing. This is the opposite of feedforward connections, called feedback pathways, and suggests that bottom-up and top-down processing both co-exist to facilitate information processing (Gilbert and Li 2013). This is supported by past research, which suggests top-down processing, guided by the alpha/beta band, works with bottom-up processing, regulated by the gamma band, in order to coordinate neuronal communication (Fries, 2008). This signal contains information to guide visual processing and create a stable image of nearby objects, despite the constant movement of saccades. In support of the discussion about the dorsal pathway as the area for visual information, feedforward connection starts primarily in visual cortex V1, and extends into the dorsal pathway, V2, V4, MT, and IT (Gilbert and Li 2013).

The relationship between physiologically facilitated depth cues (like binocular disparity and motion parallax), the spike rates occurring neuronally within the MT, and how visual information processing facilitates the vivid sensation of depth is still unanswered. In our recent models of MePMoS and NHT, the spike rates is conveyed to the gamma wave phases, then to the alpha phases (Arisaka, 2022).

## 2 Results

In this study, 3D space was modeled by a physical experimental set-up where the participant’s understanding of space was determined by the time it took them to react to stimuli at various quantifiable occurrences. These stimuli were flashing light-emitting diodes at varying distances of depth in front of the participant. Three groups of experiments were performed, an initial experiment testing various physical and procedural parameters of this investigation (2.1-2.4), a larger version of the first set-up scaled from 2.5 m in maximum depth to 5.0m of maximum depth (2.5), and an evolution of the initial concept by expanding the possible horizontal polar eccentricities where the stimuli would flash (2.6).

### 2.1 Covert versus Overt Attention

It is predicted that covert and overt attention will be similar in a linear shape, intercept and slope. When comparing the positive and negative slopes of the covert vs overt attention protocols, the positive slope creates a delay roughly 60 m/s longer than that of the negative (Figure 1). However, this is due to the nature of linearity. The negative slope of covert attention was −52.2 ± 4.3 ms/m while that of overt attention was −45.4 ± 7.7 ms/m (Table 1), we see the same pattern with the positive slopes: 69.7 ± 2.1 ms/m for overt and 67.83 ± 2.3 ms/m for covert. This coincides with our hypothesis that covert and overt attention have roughly the same reaction time, with minor deviations. Similarly, when taking a look at the intercepts for positive covert and overt attention, which were 309.0 ± 3.0 and 318.6 ± 3.0 ms/m respectively, we see that the difference in reaction time between them is minimal. The values of the reduced after the center cue for the covert and overt attention protocols were less than one, (0.62 and 0.49 respectively) suggesting that there is a strong correlation between reaction time and distances before the center cue. Despite the reduced 2of covert versus overt data after the center cue having a larger minimum chi values than the points previous to the cue, the R^2^ values for both positive and negative data points were still found to be close to one, suggesting that much of the variation in the data was accounted for by the distance variable.

**Figure 1.**
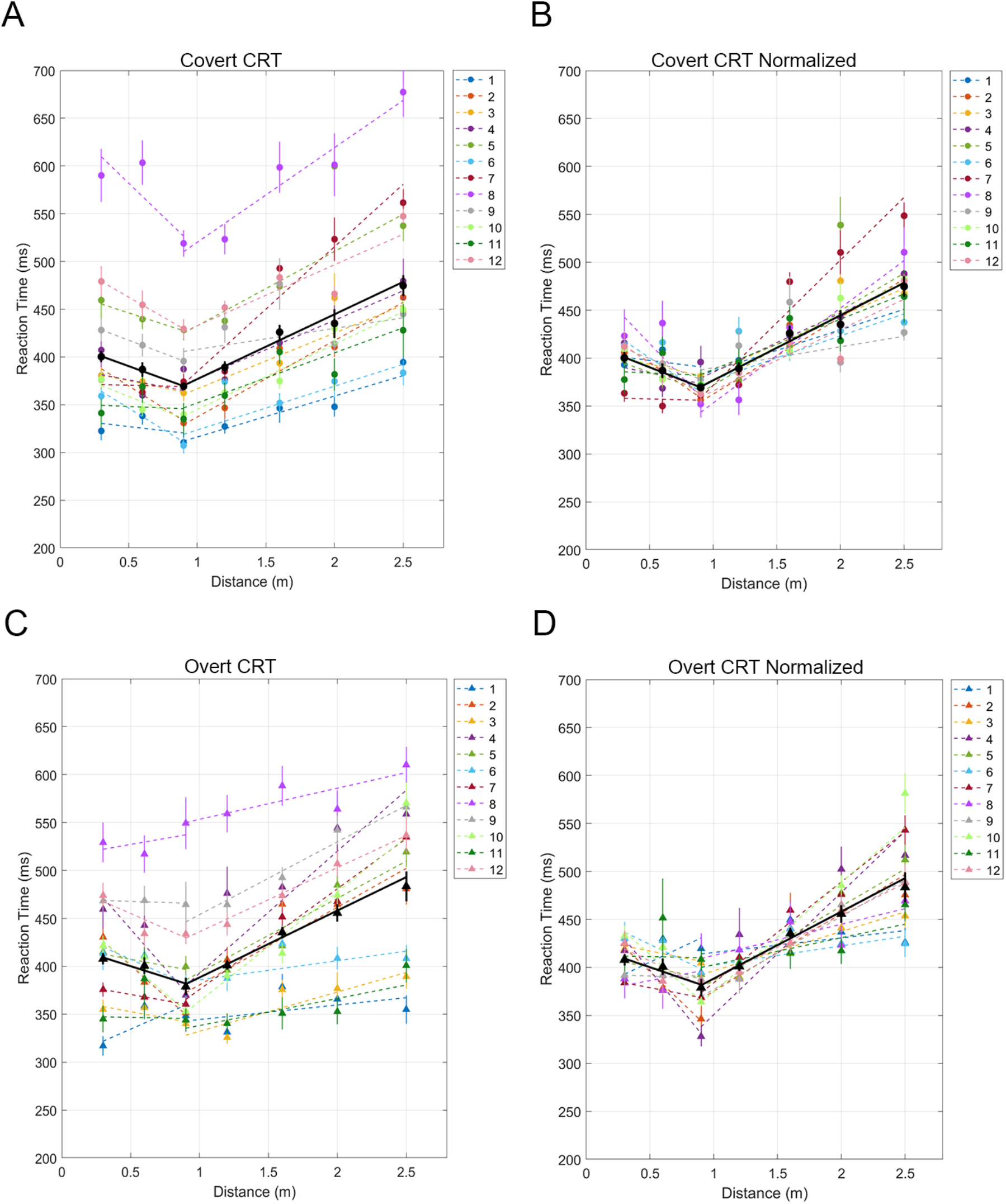
Overt and Covert Attention of Space with Stimuli at Varying Distances of Depth. Results of covert and overt attention, with the aggregation of the subjects’ data in each set denoted in black. In covert attention, the V-shape pattern is much steeper than that of overt attention. (A) Results of individual subjects observing space under covert attention. (B) The result of normalizing the individual and aggregate plots of observing space under covert attention. (C) Results of individual subjects observing space under overt attention. (D) The result of normalizing the individual and aggregate plots of observing space under overt attention.

**Table 1.**
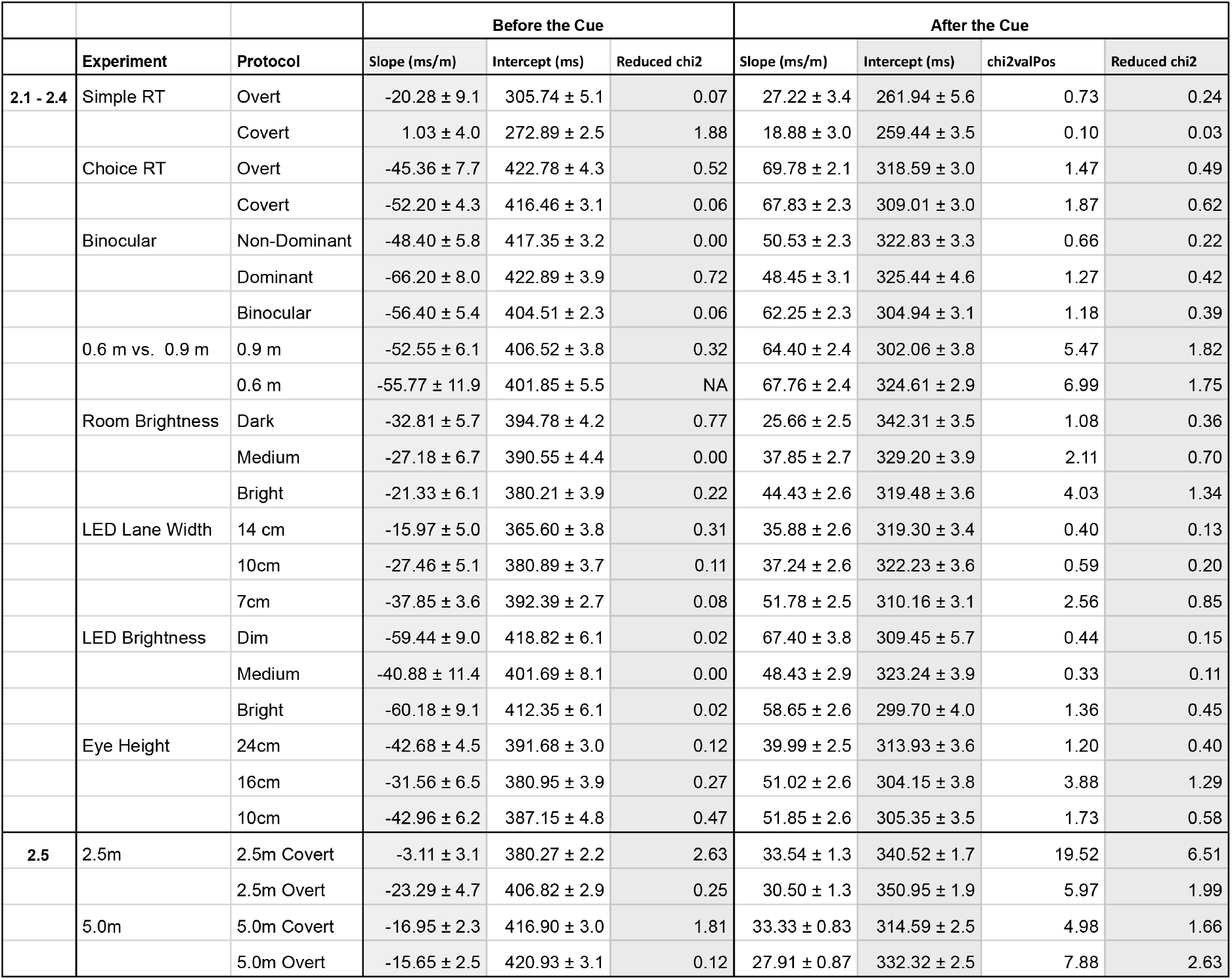
This table depicts the fit parameters of the data from 2.1-2.5 aggregated from all of the participant’s responses.

|  |  |  | Before the Cue |  |  | After the Cue |  |  |  |
| --- | --- | --- | --- | --- | --- | --- | --- | --- | --- |
|  | Experiment | Protocol | Slope (ms/m) | Intercept (ms) | Reduced chi2 | Slope (ms/m) | Intercept (ms) | chi2valPos | Reduced chi2 |
| 2.1 - 2.4 | Simple RT | Overt | -20.28 ± 9.1 | 305.74 ± 5.1 | 0.07 | 27.22 ± 3.4 | 261.94 ± 5.6 | 0.73 | 0.24 |
|  |  | Covert | 1.03 ± 4.0 | 272.89 ± 2.5 | 1.88 | 18.88 ± 3.0 | 259.44 ± 3.5 | 0.10 | 0.03 |
|  | Choice RT | Overt | -45.36 ± 7.7 | 422.78 ± 4.3 | 0.52 | 69.78 ± 2.1 | 318.59 ± 3.0 | 1.47 | 0.49 |
|  |  | Covert | -52.20 ± 4.3 | 416.46 ± 3.1 | 0.06 | 67.83 ± 2.3 | 309.01 ± 3.0 | 1.87 | 0.62 |
|  | Binocular | Non-Dominant | -48.40 ± 5.8 | 417.35 ± 3.2 | 0.00 | 50.53 ± 2.3 | 322.83 ± 3.3 | 0.66 | 0.22 |
|  |  | Dominant | -66.20 ± 8.0 | 422.89 ± 3.9 | 0.72 | 48.45 ± 3.1 | 325.44 ± 4.6 | 1.27 | 0.42 |
|  |  | Binocular | -56.40 ± 5.4 | 404.51 ± 2.3 | 0.06 | 62.25 ± 2.3 | 304.94 ± 3.1 | 1.18 | 0.39 |
|  | 0.6 m vs. 0.9 m | 0.9 m | -52.55 ± 6.1 | 406.52 ± 3.8 | 0.32 | 64.40 ± 2.4 | 302.06 ± 3.8 | 5.47 | 1.82 |
|  |  | 0.6 m | -55.77 ± 11.9 | 401.85 ± 5.5 | NA | 67.76 ± 2.4 | 324.61 ± 2.9 | 6.99 | 1.75 |
|  | Room Brightness | Dark | -32.81 ± 5.7 | 394.78 ± 4.2 | 0.77 | 25.66 ± 2.5 | 342.31 ± 3.5 | 1.08 | 0.36 |
|  |  | Medium | -27.18 ± 6.7 | 390.55 ± 4.4 | 0.00 | 37.85 ± 2.7 | 329.20 ± 3.9 | 2.11 | 0.70 |
|  |  | Bright | -21.33 ± 6.1 | 380.21 ± 3.9 | 0.22 | 44.43 ± 2.6 | 319.48 ± 3.6 | 4.03 | 1.34 |
|  | LED Lane Width | 14 cm | -15.97 ± 5.0 | 365.60 ± 3.8 | 0.31 | 35.88 ± 2.6 | 319.30 ± 3.4 | 0.40 | 0.13 |
|  |  | 10cm | -27.46 ± 5.1 | 380.89 ± 3.7 | 0.11 | 37.24 ± 2.6 | 322.23 ± 3.6 | 0.59 | 0.20 |
|  |  | 7cm | -37.85 ± 3.6 | 392.39 ± 2.7 | 0.08 | 51.78 ± 2.5 | 310.16 ± 3.1 | 2.56 | 0.85 |
|  | LED Brightness | Dim | -59.44 ± 9.0 | 418.82 ± 6.1 | 0.02 | 67.40 ± 3.8 | 309.45 ± 5.7 | 0.44 | 0.15 |
|  |  | Medium | -40.88 ± 11.4 | 401.69 ± 8.1 | 0.00 | 48.43 ± 2.9 | 323.24 ± 3.9 | 0.33 | 0.11 |
|  |  | Bright | -60.18 ± 9.1 | 412.35 ± 6.1 | 0.02 | 58.65 ± 2.6 | 299.70 ± 4.0 | 1.36 | 0.45 |
|  | Eye Height | 24cm | -42.68 ± 4.5 | 391.68 ± 3.0 | 0.12 | 39.99 ± 2.5 | 313.93 ± 3.6 | 1.20 | 0.40 |
|  |  | 16cm | -31.56 ± 6.5 | 380.95 ± 3.9 | 0.27 | 51.02 ± 2.6 | 304.15 ± 3.8 | 3.88 | 1.29 |
|  |  | 10cm | -42.96 ± 6.2 | 387.15 ± 4.8 | 0.47 | 51.85 ± 2.6 | 305.35 ± 3.5 | 1.73 | 0.58 |
| 2.5 | 2.5m | 2.5m Covert | -3.11 ± 3.1 | 380.27 ± 2.2 | 2.63 | 33.54 ± 1.3 | 340.52 ± 1.7 | 19.52 | 6.51 |
|  |  | 2.5m Overt | -23.29 ± 4.7 | 406.82 ± 2.9 | 0.25 | 30.50 ± 1.3 | 350.95 ± 1.9 | 5.97 | 1.99 |
|  | 5.0m | 5.0m Covert | -16.95 ± 2.3 | 416.90 ± 3.0 | 1.81 | 33.33 ± 0.83 | 314.59 ± 2.5 | 4.98 | 1.66 |
|  |  | 5.0m Overt | -15.65 ± 2.5 | 420.93 ± 3.1 | 0.12 | 27.91 ± 0.87 | 332.32 ± 2.5 | 7.88 | 2.63 |

The aggregate data regarding the trials with overt attention displayed marginally slower reaction times than that of covert attention (Figure 1). However, we note that despite this slight difference in reaction times produced between the varying attention, for a single subject, the V shape patterns are almost identical, as we had anticipated for covert and overt attention (Figure 1).

### 2.2 Depth is Converted to Time

These next experiments compared the difference between the previously conducted choice reaction time experiment, where the participant indicated whether one or two LED’s appeared, and a simple reaction time experiment, where only a single stimulus could appear at each point. Looking at (A) from Figure 2, the positive and negative slopes are significantly less than the positive and negative slopes of (B) from Figure 2. A numerical comparison shows that the positive covert and overt slopes of the simple reaction time experiment offer 27.22 ± 3.4 m/ms and 18.88 ± 3.0 m/ms respectively, while the choice reaction time experiments positive covert and overt slopes increase to 69.78 ± 2.1 m/ms and 67.83 ± 2.3 m/ms respectively. This apparent flattening of the slopes with the simple reaction time experiment indicates that the reaction time delay produced as a function of distance is increased with conscious decision-making as the mechanism for responding to the reaction stimulus.

**Figure 2.**
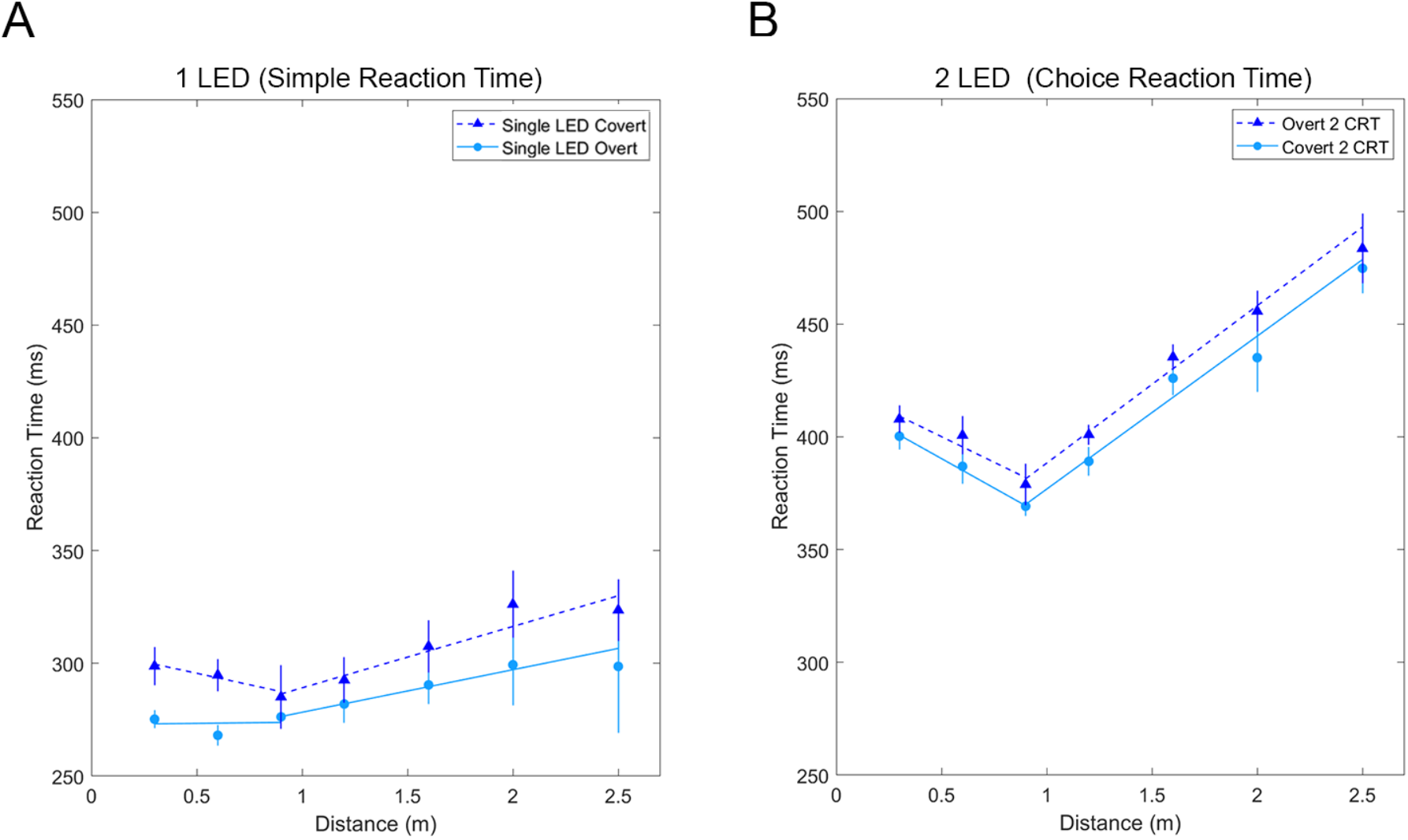
Simple Reaction Time versus Choice Reaction Time under Overt and Covert Attention. (A) Results of data taken in reaction to a single LED flashing randomly on either side of the LED strip. These trials are considered to be simple reaction time, and because of this we note that the resulting patterns are relatively flat. (B) Results of trials taken with a 0.9 m center cue using both covert and overt attention. While overt attention gives a steeper slope, we note that both meet at the same vertex, and that the differences in the slopes are minimal.

### 2.3 Monocular and Binocular Data and the Impact of Different Focusing Distances

Both the binocular and monocular data show significant slopes (Figure 3). After the center cue, the binocular data had the greater slope, with the more distal points demonstrating an average reaction time 62.2 ± 2.3 ms slower than the reaction time of a meter closer, shown as the slope (with a unit of ms/m) in the relevant graph. The singular dominant and non-dominant eye protocols had slopes of 48.5 ± 3.1 ms/m and 50.5 ± 2.2 ms/m, respectively (Table 1). Before the center cue, the dominant eye had the largest difference in reaction time, with an average difference of −66.2 ± 8.0 ms/m for lower distances, while the binocular and non-dominant eye protocols showed on average a −56.4 ± 5.4 ms/m and −48.4 ± 5.8 ms/m lower reaction time for a larger distance, respectively. This data supports the hypothesis that binocular vision is not necessarily needed to perceive depth as each individual eye showed significant slopes on their own.

**Figure 3.**
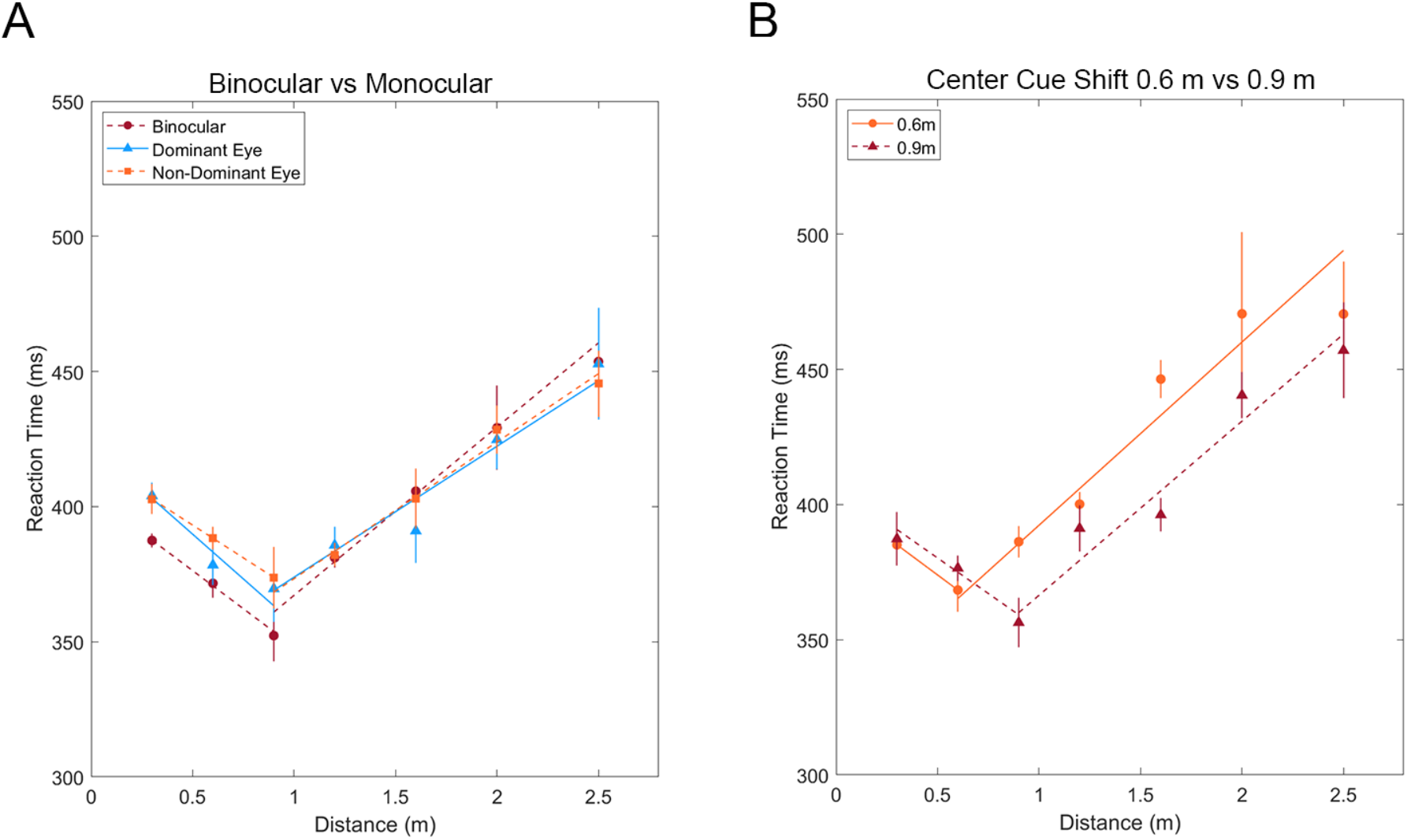
Binocular vs Monocular Viewing of Depth and Varying Center Cue Distances. (A) Data of trials taken with both eyes, the dominant eye, and the non-dominant eye (n = 8). While binocular vision exhibits the shallowest V-shape pattern, both data sets of single eyes show that a single eye is enough to perceive depth. (B) Data of trials taken with a center cue at the standard 0.9 m, and at 0.6 m. From this, we see that the distance of fastest reaction time adjusts accordingly.

For trials conducted with both eyes, the cue flashing at 0.9 m, where visual gaze rested throughout trials, resulted in the fastest reaction time, with an average of 304.9 ± 3.1 ms. The participants’ reaction times then increased from this cue position with increasing distance in a linear manner. From 0.9 m, extending to either 0.3 m, or 0.6 m away from the focus of vision, the reaction time to flashing stimuli would increase.

From the trials conducted with varying center cues (0.6 m from the participant vs. 0.9 m from the participant), the findings displayed a change in the position of the kink in the graph, or distance with the lowest reaction time, that corresponded to the different cues (Figure 3). For the trials in which the participants’ gaze was held at 0.6 m away, the fastest average reaction time across the trials was 324.61 ± 2.9 ms, and at the 0.6 m center cue. Whereas the fastest average reaction time across trials for the 0.9m center cue comparative experiment was 302.06 ± 3.8 ms, but at the 0.9 m center cue’s position. The reaction times in both cases increased linearly, and for distances up to 2.5 m, or 2 m from the observer, resulted in the greatest total reaction times.

We hypothesized that the various reaction times when plotted would create a V-shape pattern, with its vertex at the location of the center cue, and that shifting the center cue would have no effect on creating this pattern. This was supported by our data, as the 0.6 m and 0.9 m center cue protocol had similar slopes for both the before center cue and after center cue data, with the slopes being −55.8 ± 11.9 ms/m and −52.5 ± 6.1 ms/m for the closer data points, and 67.8 ± 2.4 ms/m and 64.4 ± 2.4 ms/m for the farther points, respectively (Table 1).

With the exception of the single LED trials’ (simple choice) flatter slope, we found that all trials exhibited the same pattern, with the fastest reaction time being at the center cue, and reaction times increasing linearly with distance away from this point (Figure 3). Furthermore, this was the same pattern even when the center cue was shifted from the standard location to a closer point of focus.

### 2.4 Control of Various Parameters

In order to fully understand the V-shape pattern appearing consistently across the previously discussed experimental protocol, the physical qualities of the experiment were tested to see if they altered the results. These physical qualities were the ambient light of the room the subject was in, the brightness of the stimulus LED the subject responded to, the width between the LED strips and the height of the subject’s eyes.

The LED stimulus was controlled using the Arduino UNO and its internal unit for luminosity. We set this unit for the LEDs and center cue at 219.5 Mcd for the dim trials, and respectively set it at 239.1 and 258.6 Mcd’s, for the Medium and Bright trials. Room brightness was tested as each subject changed the level of ambient light in the space they performed the experiment, with “bright room” representing full, indirect natural daylight, “medium room” meaning indirect daylight in the later afternoon and “dark room” meaning a room with no daylight and dim artificial light. Lane width and eye height refer to the physical dimensions of the experimental setup used. The lane width between the LED strips and the eye height of the subject were increased incrementally and tested. All of these varying conditions produced data that held little inconsistency between the various trials. (Figure 4) This determined that these various changes in the testing parameters did not alter the consistency of the pattern regarding increased stimulus eccentricity from the center cue producing a greater delayed response from the subject.

**Figure 4.**
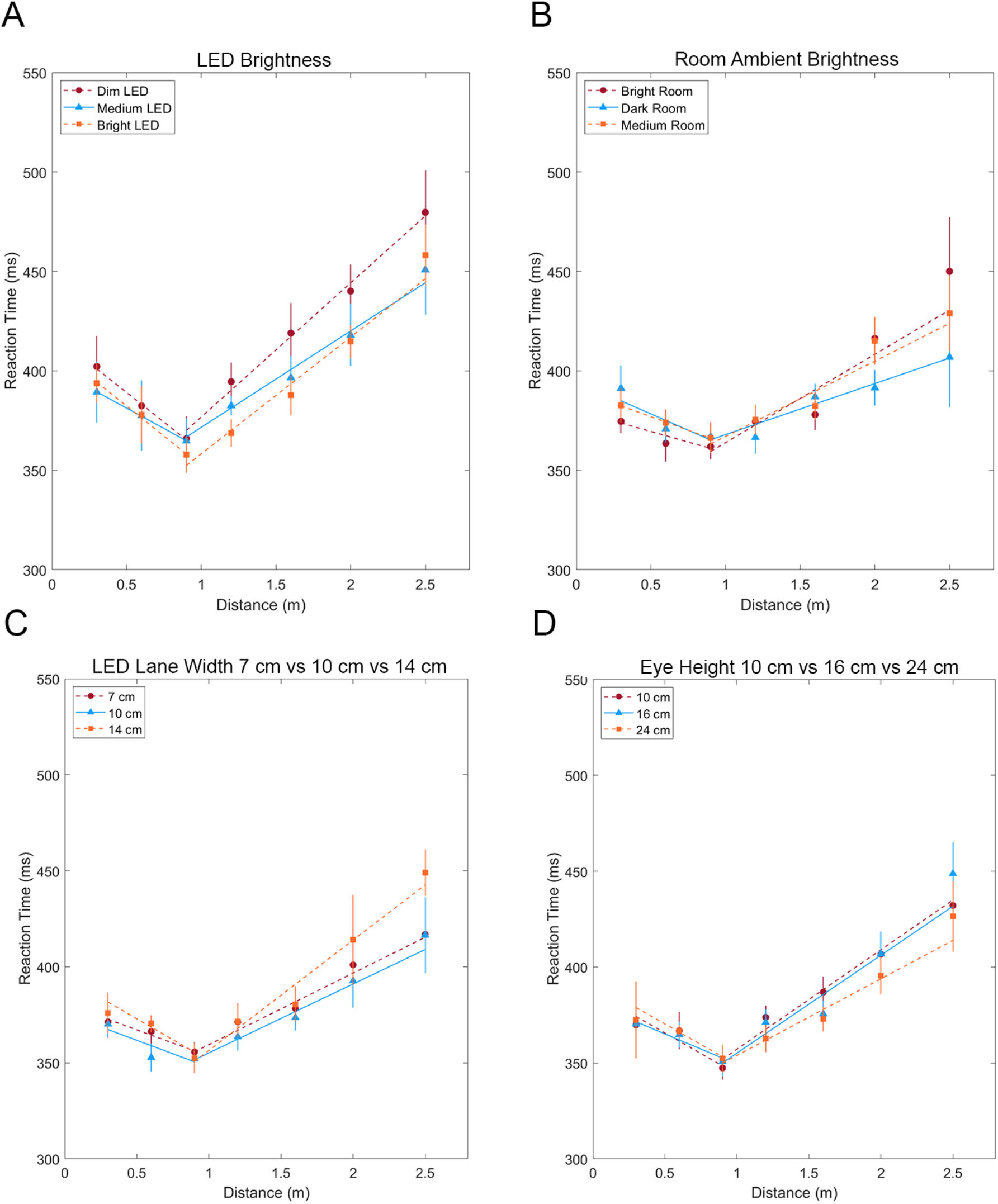
Varying the Physical Parameters of the LED Depth Experiment. (A) Results of data taken with the LEDs set at different brightnesses. Using the Arduino UNO, we set the brightness of the LEDs and center cue at 219.5 Mcd for the dim trials, and respectively set it at 239.1 and 258.6 Mcd’s, for the Medium and Bright trials. The V-shape pattern remained intact, and we conclude that any brightness works, so long as it is not too dim, nor too bright that it hurts the subject’s eyes. (B) Results of data taken in rooms with different lighting. We found that as long as there is some ambient light, we are still able to perceive depth. (C) Data of trials taken with various distances between the LED lanes. Similar to changing eye height, we saw no significant deterioration in the overall pattern. (D) Data of trials taken at different eye heights. While we were initially using a 0.10 m eye height, where the chin of the subject is directly on the same surface as the LED strip, we found that setting our eye level at even higher distances did not have much effect on the V-shape pattern.

### 2.5 Scaling Space by Increasing the Proportions of the Experimental Set-Up: 2.5 m to 5.0 m

This next experiment consisted of an identical version of our previous 2.5m Covert vs. Overt protocol, however this time, all the physical proportions were scaled up by a factor of two, and the subject was sitting, rather than prone. Simultaneously we reconstructed and reconducted our old 2.5m experiment in a sitting fashion, as all previous experimentation had been done remotely and prone. We hypothesized our V pattern would hold. We note that all positive slopes for all Covert vs Overt trials, on both setups, were around 30 ms/m, shrunken from the magnitude of the positive slopes in some of the previous experimentation. We also note that all negative slopes for all trials also shrunk in magnitude. It is possible this is due to the greater practice with the experimentation by previous subjects. It is important to note that with this change, chi-squared and reduced chi-squared values for some of the trials were slightly higher than previous experimentation, but still close to one. It is possible that this was due to a subject pool more unfamiliar with any sort of data taking or is simply due to the increased size of the experiment, and more depth into space. Regardless, it is clear our V shape remained intact, as did the linear perception of depth based on distance from the cue, even though it was not as steep.

**Figure 5.**
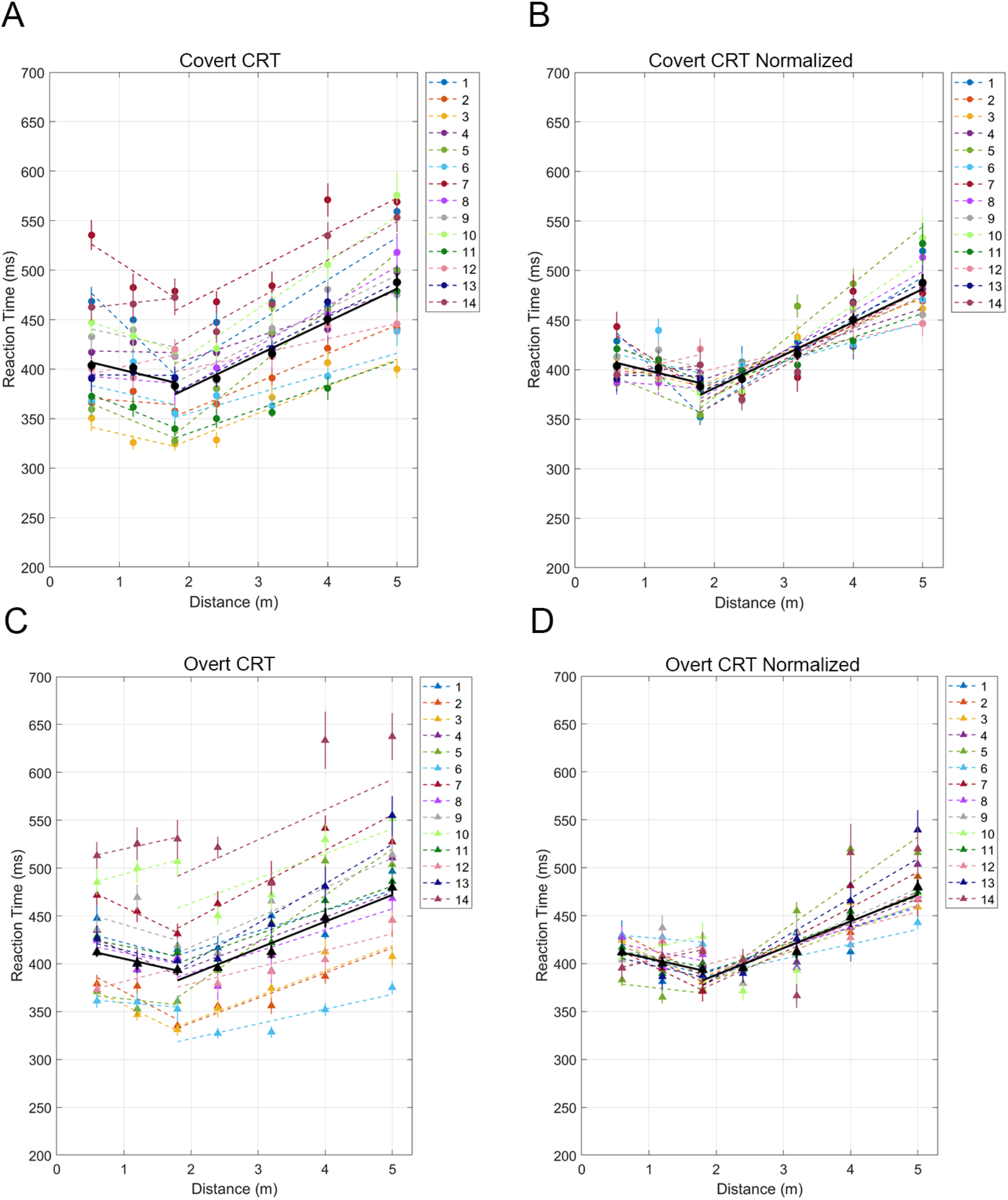
Scaled 5.0m Depth Results of covert and overt attention experiment. done on a 5.0m long setup, with the aggregation of the subjects’ data in each set denoted in black. In covert attention, the V-shape pattern is nearly equivalent to overt attention. (A) Results of individual subjects observing space under covert attention. (B) The result of normalizing the individual and aggregate plots of observing space under covert attention. (C) Results of individual subjects observing space under overt attention. (D) The result of normalizing the individual and aggregate plots of observing space under overt attention.

### 2.6 Depth and Horizontal Eccentricity as a Model for Two-Dimensional Space

The Two-Dimensional lattice setup was an extension of our previous depth protocols, with a factor of eccentricity added. In order to do this we placed multiple LED strips, horizontally at different points of depth. This would allow us to flash stimuli at our previous depths with 0°of eccentricity and then also extend to ±20°,and ±40°of eccentricity. All trials were conducted with Covert attention, and with the subject seated. Taken with N=5, these plots show strong correlations between varying eccentricities leading to a change in reaction time, and also varying depths, leading to a change in reaction time. When comparing the data plotted in respect to polar angles versus the data plotted in respect to the Cartesian-X Distance, we see possible hints of a polar coordinate system farther away from the subject (>1.8m) and hints of a Cartesian coordinate system up closer to the subject (<1.8m).

## 3 Discussion

Our results did demonstrate a negligible difference between covert and overt attention when it comes to the conscious interpretation of spatial information. It was hypothesized that covert and overt reactions would be nearly identical. Saccades occur unconsciously especially during covert attention when the eyes are supposed to be fixed at one location. When we move our eyes towards an object to determine its identity, we are exercising overt attention, with the brain comparing its predicted semantic interpretation of the object to reality based on prior experience. The positive slope’s intercepts were 318.59 ± 3.0 ms for overt attention and 309.01 ± 3.0 ms for covert attention; the reaction times for distances further away continued to be slightly slower for overt compared to covert. The positive slopes themselves were 69.78 ± 2.1 ms/m and 67.83 ± 2 for overt and covert, respectively, showing the parallels between the two attentions. The discrepancy between the two is so slight, that it demonstrates the strong indifference between the two modes of attention. Such a small discrepancy from our hypothesis could likely be attributed to subject error, or a limited subject pool.

The data regarding the monocular vision of the participants’ dominant or non-dominant eye, and binocular vision all demonstrate clear V patterns that suggest proper depth perception. As humans utilize the same pathways during monocular and binocular vision, it was predicted that they would produce similar results in regard to depth perception (O’Keefe 1978).

There are many factors that could impact the reproduction of the slope values of each participant over the different experiments we have conducted. Prior to the 2.5m established system and the 5m setup, the participants were becoming increasingly familiar with the experiment, and the environment itself for most participants was known to them, as participants set up their experiments at home. These retrieval processes contribute to the top-down processing described earlier and may very well enhance the prediction accuracy of participants over time, altering the slopes. Along with this, all such data was taken in a prone position, whereas once there was the established 2.5m system and 5m scaled system and the 2D Lattice system, all data was taken in an upright position. We note that the size of the room that the experiment is conducted in, and objects in it, may also play a role in the top-down processing system. More testing would be necessary.

Based on the results of this study, depth perception was observed in multiple conditions, including covert and overt attention, binocular and monocular vision, at different levels of room brightness, and at differing center cues. The linearity of the increasing reaction times extending from the center cue and the lowest reaction times observed at the center cue, the focus point of our visual frame, points to an internal remapping of this external space that acknowledges the distances between landmarks. Further research can be explored in the processes with monocular depth perception as data suggested binocular vision is not necessary to properly perceive depth. Further research can also be done with quantifying the amount of ambient light needed for depth perception, as our data suggests depth perception weakens in the dark compared to light conditions.

From our specific control parameters, such as a 0.07 m distance between each LED strip, an eye height of 0.10 m, proportional to the subjects laying their chin on the same surface, a standard LED brightness of 25 by the Arduino unit, and the ambient light of the room, we varied these four external factors and examined how changing such parameters would affect our results. In Figure 4, while there are variations of reaction time, overall these variations are minimal. The same linear pattern was achieved across all data sets, with a linearly dependent relationship between 3D space and time.

## 4 Methods and Materials

### 4.1 Participants

Participants were drawn from the undergraduate and graduate UCLA student population in accordance with approved procedures from the Institutional Review Board (IRB # 19-001472). Due to the COVID-19 pandemic, data-taking was conducted remotely with internal members from the summer of 2020 until the summer of 2021. After which, standardized experimental setups were developed in-lab and external participants were recruited for additional, professional data-taking. These groups’ data were assessed separately for consistency, then combined in an aggregate analysis as shown in the Results.

No subjects nor participants had any history of serious visual abnormalities, and vision correction was required in any case of the subject having visual abnormalities. The 5m scaled setup, along with a singular and established 2.5m system, was then built and operated on campus with the same previous covert versus overt protocol as conducted remotely.

### 4.2 Ocular Dominance Determination

Participants conducted a standardized eye preference test prior to data taking to determine their dominant and non-dominant eye. Each subject was instructed to fully extend both arms and create a triangular opening using opposing thumbs and index fingers. The triangle shape was then centered on a distant object. If the object remains centered when the left eye is closed, the participant is considered to be right eye dominant. If the object is out of frame with the left eye closed, the participant is considered to be left eye dominant.

### 4.3 Setup

The 2.5m setup consists of an Uno Rev3 Controller Board, breadboard, pushbuttons, jumper wires, and 10 kΩ resistors from Arduino LLC, located at 10 St. James Avenue, Boston, Massachusetts, 02116, USA. Subjects also used a 5 m WS2812B 300 pixel light emitting diode (LED) strip from ALITOVE Electronic Technology Co., Ltd., located at 4B 7 Danyuan 12 Dong, Minzhi Street, Longhua District, Shenzhen, Guangdong, 518000, China. To power and upload code into the LED strip, participants used a computer with the Arduino IDE. The 5m setup is identical apart from the additional light strip, and a headrest for the participant, which was also included on the established in-lab 2.5m system.

Two pushbuttons with 10 kΩ resistors, were placed on the breadboard and connected with jumper wires to the Arduino Uno Rev3 controller board (Figure 6). Each pushbutton was wired in series with a resistor. The Arduino was then connected to a computer which could upload code and provide power. The LED strip was connected to the Arduino and the breadboard via a three pin Japan Solderless Terminal SM series connector, with the green wire for data input, the black wire to ground, and the red wire to a 5V power supply from the Arduino. Each LED on the strip was individually addressable and programmable with specific color and animation. The 5 meter LED strip was taped down to a level surface and bent at the halfway point to create two parallel 2.5 m strips (Figure 6). In the case of the 5m setup, each strip was harnessed parallel to a level surface, and ran their full length; there was no bend in either strip. These individual strips were then separated by 7 cm; 14 cm for the 5m setup, as it is difficult to differentiate the two strips if the separation is too minimal and also difficult to pay attention to both strips at once if the separation is too large. Each subject ensured that at the end of their LED strip there was a solid background, either a wall or a large board. An object of the subject’s choice (typically black tape) was placed at the center cue, the position at which the subject would focus their gaze (covert attention), or return their gaze after reacting to an LED flash (overt attention). The surrounding light was relatively dim, but not completely dark as to block out all other light, so LED flashes could be perceived more clearly. (Figure 7).

**Figure 6.**
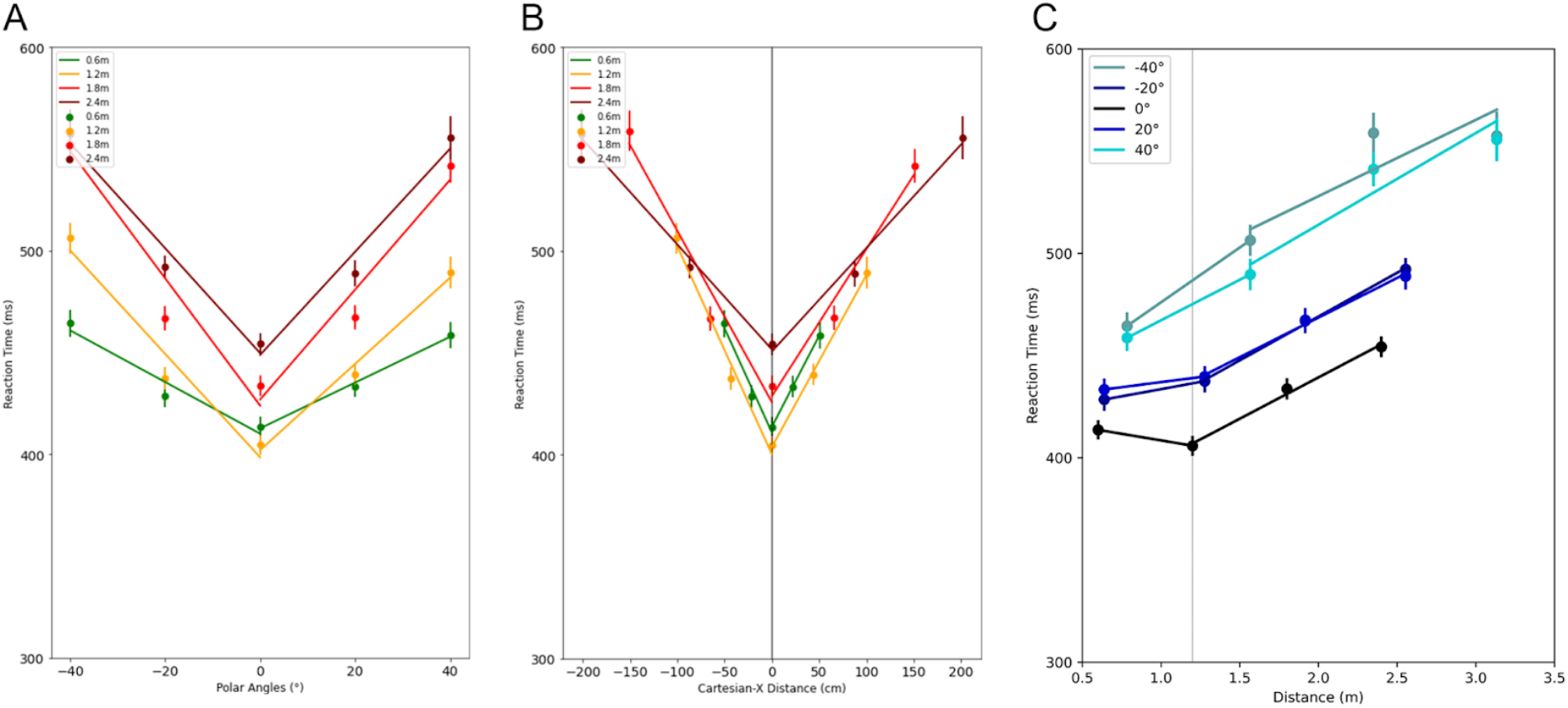
Two-Dimensional Lattice, Reaction Time as a function of Polar Angles and Depth. These plots represent the results of recording the reaction times of participants as they respond to stimulus LED flashes at various locations as determined by a depth (0.6m, 1.2m, 1.8m, 2.4m) and an angle of eccentricity (±40°, ±20°, and 0°) (A) Reaction time (ms) in relation to the polar angle (°) where stimuli had flashed. (B) Reaction time (ms) in relation to the distance between the participant and stimulus by the x-component of a cartesian representation of space (C) Reaction time (ms) in relation to the radial distance of the stimulus from the subject. The 0° eccentricity has the center cue of this experiment marked at 1.2m from the subject.

**Figure 6.**
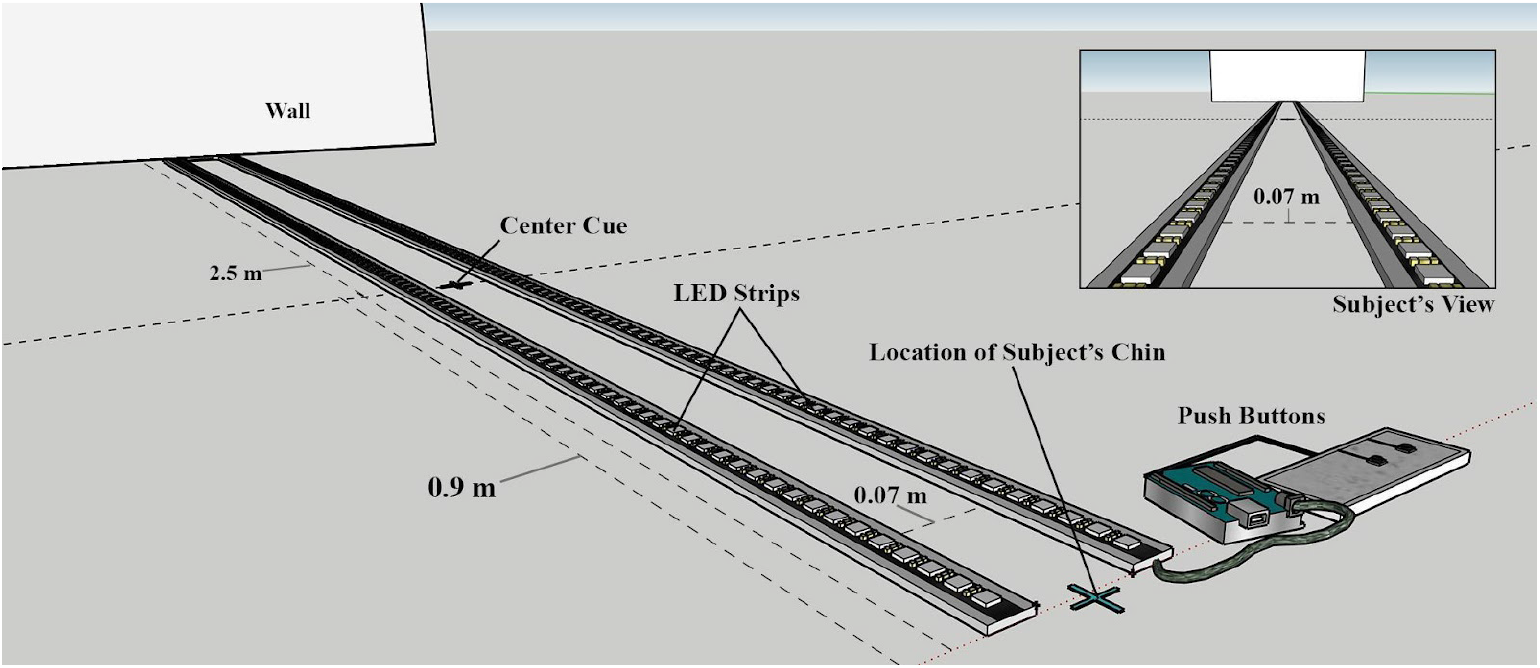
An overall view of the entire setup. A subject’s point of view of the experimental setup is depicted as well. Subjects would look straight ahead, focusing their attention (covert) or returning their gaze (overt) to the center cue (dotted line). To increase the subject’s concentration on the center cue, an object of their choice was placed at the center cue mark, such as a sticky note or solid object. The distance between the two lanes is 0.07 m; increasing the distance further would make it more difficult to perceive both lanes together. The LED strip length is 2.5 m as it becomes more difficult to perceive anything past this distance. The subject would rest two fingers on the push buttons and press the buttons according to what they perceived: two LEDs flashing or a single LED on either lane.

**Figure 7.**
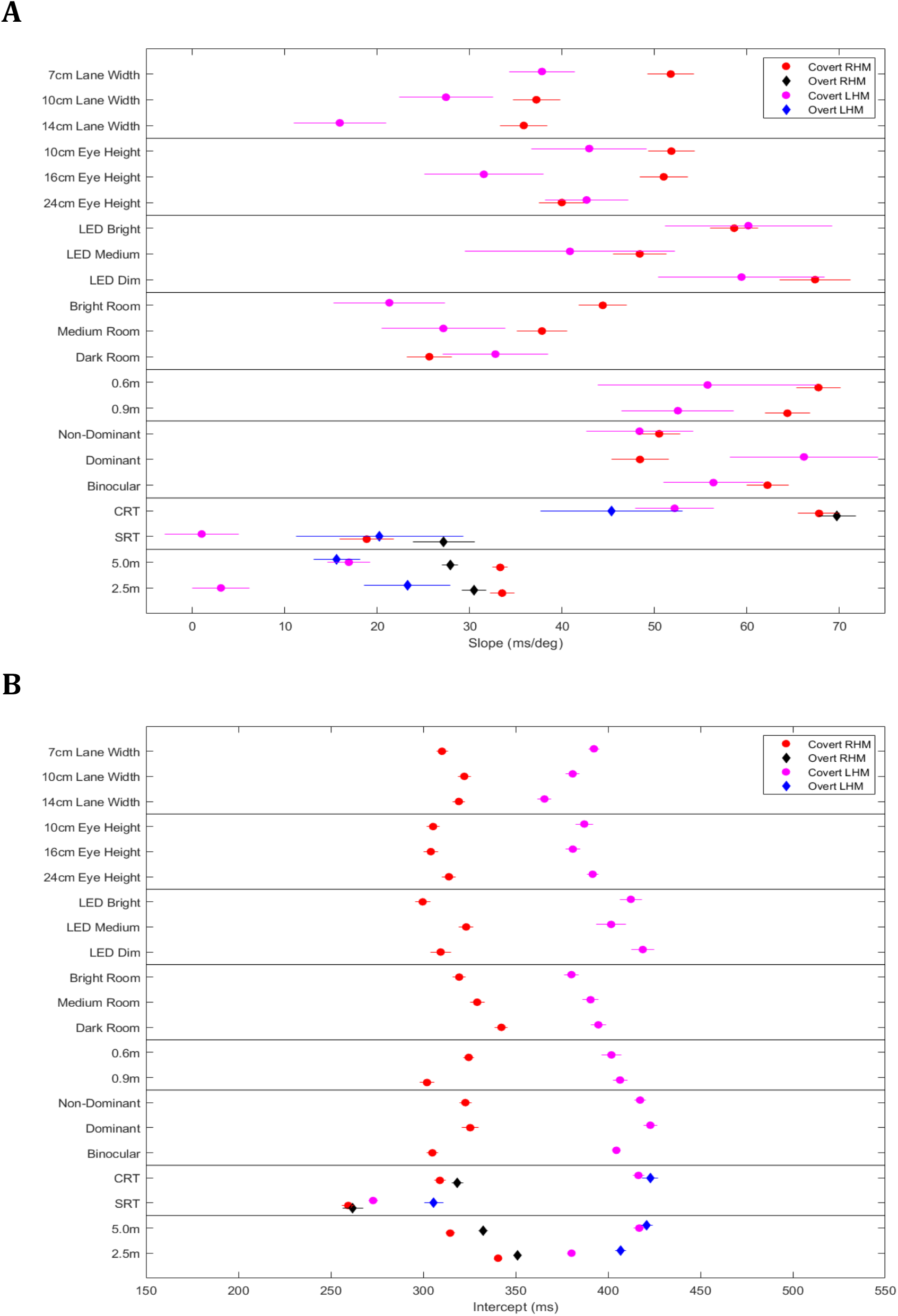
Slope-Intercept Analysis Across all Experiments. (A) This figure compares the slopes (ms/m) of reaction time vs distance across the 8 data sets. (B) This figure compares the intercepts of reaction time (ms) across the 8 data sets.

**Figure 7.**
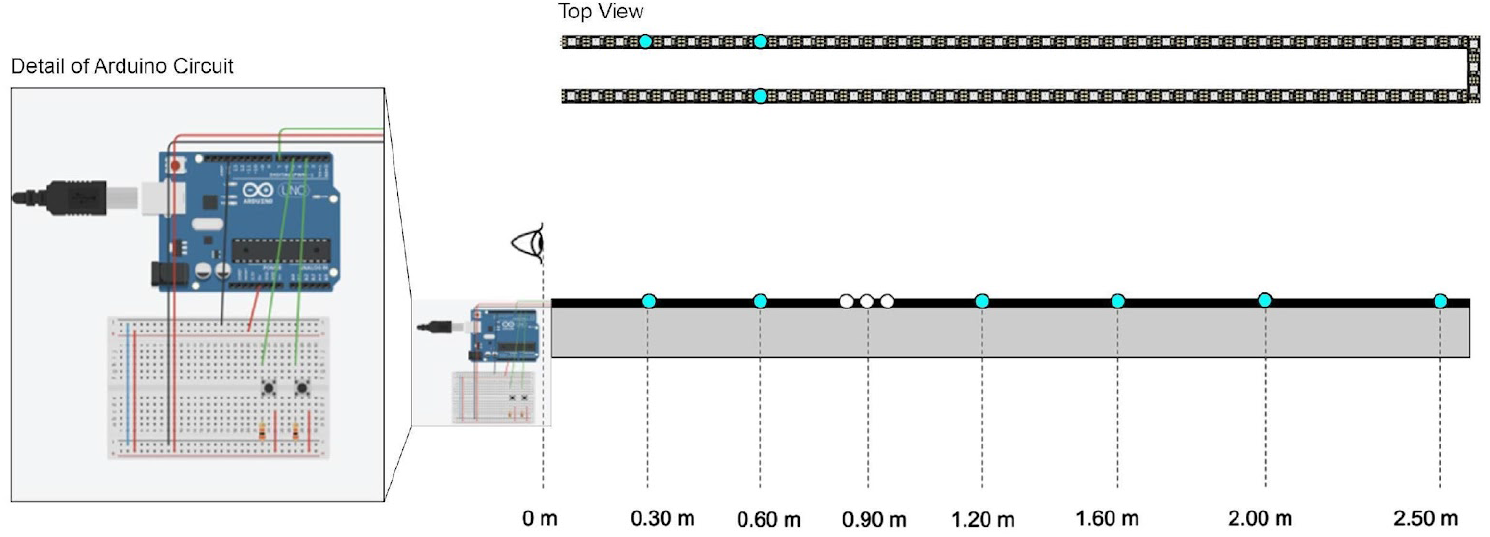
An overall view of the entire setup. An aerial view of the experimental setup is depicted as well. Subjects would look straight ahead, focusing their attention (covert) or returning their gaze (overt) to the center cue (dotted line). To increase the subject’s concentration on the center cue, an object of their choice was placed at the center cue mark, such as a sticky note or solid object. The distance between the two lanes is 0.07 m; increasing the distance further would make it harder to perceive both lanes together. The LED strip length is 2.5 m as it becomes more difficult to perceive anything past this distance. The subject would rest two fingers on the push buttons and press the buttons according to what they perceived: two LEDs flashing or a single LED on either lane.

**Figure 8.**
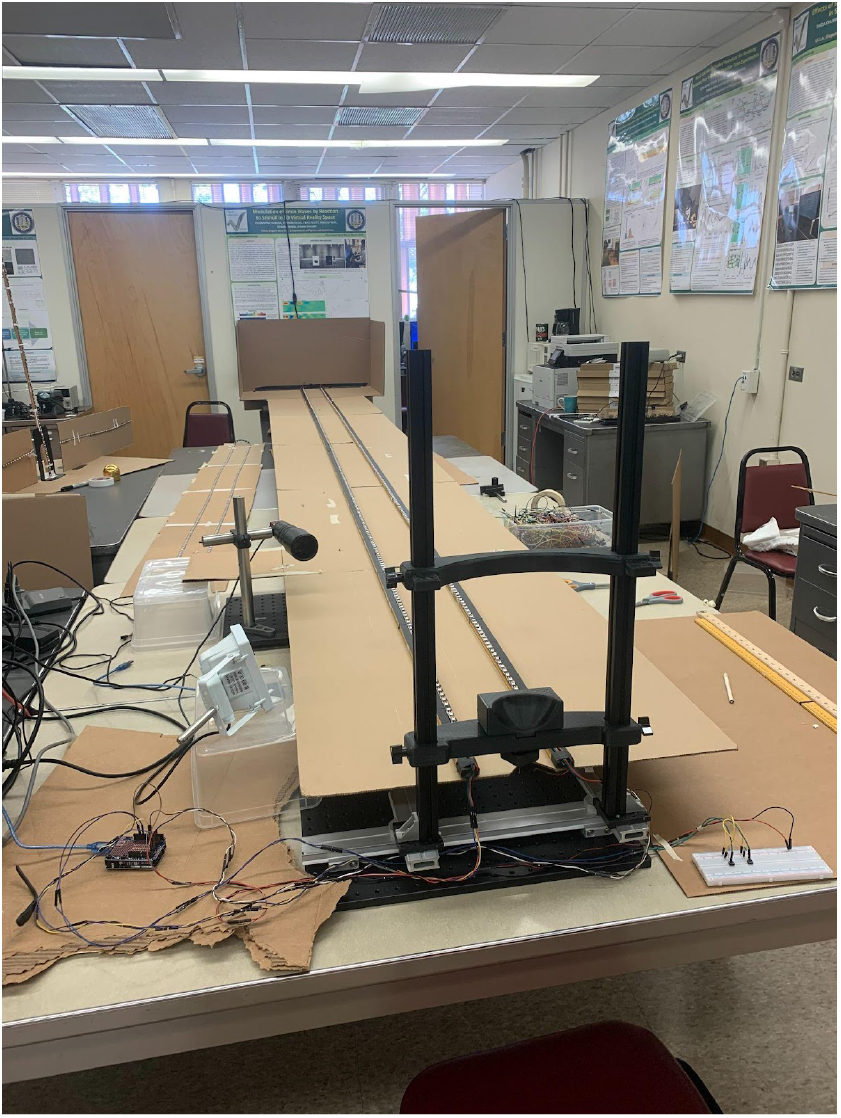
The 5m LED Depth Setup used at UCLA Knudsen Hall.

**Figure 9.**
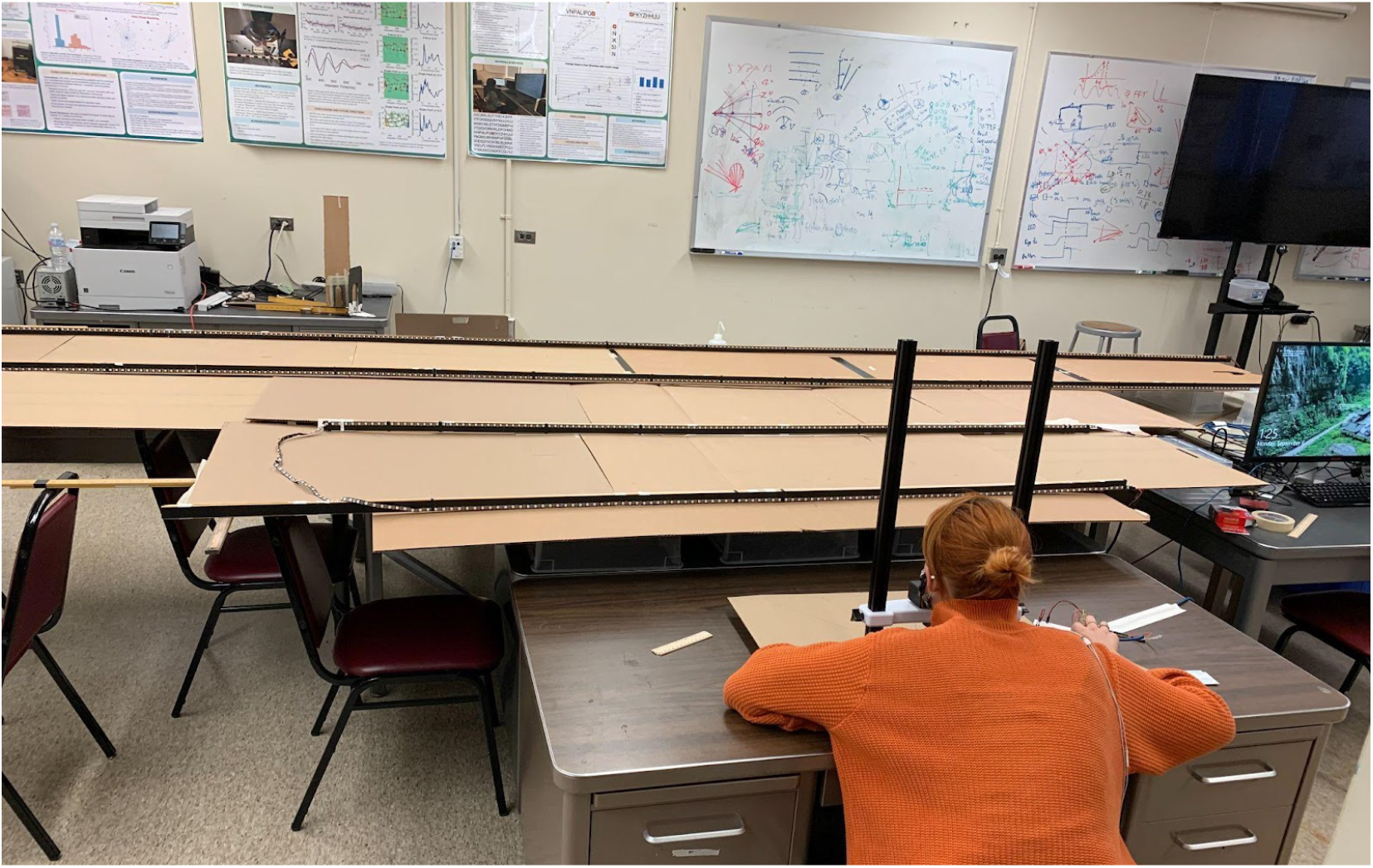
The Two-Dimensional Lattice with depths at 0.6 m, 1.2 m, 1.8 m, 2.4 m and points of horizontal eccentricity at ±40°, ±20°, and 0°.

### 4.4 Procedure

For all experiments, participants took multiple sets of data within various specified parameters. Each data set consisted of at least 200 individual successful trials.

The first experiment required three sets of data, with a center cue at 0.9 m. In the first data set, the participants used both eyes (binocular vision), the latter two sets of data were then conducted with monocular vision using each dominant and non-dominant eye. This experiment was conducted using covert attention, where the subject’s gaze was held as the center cue for the duration of the experiment, and took place on the 2.5m setups.

The second experiment was conducted using two of these data-sets of 200 trials to compare the effect of different center cue positions on our V shape. The first set was taken with the center cue at 0.6 m, whereas the second set was taken with the center cue at 0.9 m. Each set of data was conducted with covert attention, just like in the monocular vs binocular experiment prior, and took place on the 2.5m setups.

The third experiment, inspecting the differences between covert versus overt attention, was again conducted by taking two data-sets. The cue position would return to the. 9m point for both sets of data, and they would be taken on the 2.5m systems. The first set was taken with the subject’s focus at the center cue at all times throughout the two hundred trials (covert attention), and the second set was at a 0.9 m center cue with the subjects being able to move their eyes freely, while returning their focus to the center cue when it flashed between each individual trial (overt attention).

The fourth experiment tested if changing various parameters, such as ambient room brightness, LED brightness, separation of the LED strips, and height of the participants’ heads in relation to the surface of the LED strips would impact the pattern observed from previous data. The first of the parameters tested was a bright versus medium bright versus darkroom experiment with a center cue at 0.9 m. For this procedure, participants were instructed in the first set to simulate a bright setting with daylight to dampen the effects of the bright LED flashes. The second set of data was to be conducted without this bright direct natural light, with more ambient lighting, and the third, in a dark room, with no natural light, meant to contrast heavily with the bright LED flashes. The second parameter regarding LED brightness was done with three data sets with the LEDs set to a particular brightness for each set. The third parameter testing lane width tested three individual separations of 7, 10, and 14cm. The fourth parameter tested 3 individual subject head heights of 10, 16, and 24 cm. All of these data sets were taken with covert attention, and on the individual 2.5m setups.

The fifth experiment tested if changing various parameters of the physical set-up impacted the pattern observed from prior data. These physical changes included extending the setup to 5m, raising the headrest, and spreading the lanes. Participants were instructed to take two sets of data, one conducted with covert attention, and another with overt, and then again on the 2.5m setup. These were all conducted in a seated position, all previous experimentation was conducted in a prone position.

The sixth experiment was meant to test if increasing eccentricities, tied with increasing depths, had a predictable trend for reaction times. This setup included four horizontally placed LED strips at depths of 0.6m, 1.2m, 1.8m, and 2.4m. Participants were instructed to record four data sets of 200 trials, each set identical, with covert attention, and with the LEDs flashing at any of these depths and eccentricities of ±40°, ±20°, and 0°. This was also conducted in a seated position. The four data sets were then compiled into one for each subject.

For all experiments, the flashing LED color was set to blue, and the center cue consisted of three adjacent white LEDs. The participants rested their chins approximately on the same surface that the LED strip was secured on, leaving their eyes 10 cm above the LED strip (the distance from the participant’s eyes to the LED strip surface). The experiment consisted of two conditions, where a single LED flashed on either the left or right side of the parallel LED strips, or two adjacent LEDs flashed, one on each parallel light strip. The participants were instructed to press the left pushbutton in the case of a single LED flash, and the right push-button if two LEDs flashed. The LED strip was programmed to flash randomly within a set 300 to 800 milliseconds inter-stimulus interval. The distance and number of LEDs flashed were randomized to prevent the participants from expecting a certain flash at a predetermined location. Subjects were instructed to react to the stimulus immediately by pressing the correct button corresponding to the number of LEDs flashed. For single eye data collection, the participants’ non-dominant hand was utilized to cover one eye, while the dominant hand was used to respond to LED flashes. For binocular vision, the participants’ dominant hand was used to press the push buttons. All trials involved the index finger on the pushbutton corresponding to one LED and the middle finger on the pushbutton corresponding to two LEDs. The flashes were spaced throughout distances of 0.30, 0.60, 0.90, 1.20, 1.60, 2.00, and 2.50 meters from the subject on the 2.5m setup, and at distances of 0.60, 1.20, 1.80, 2.40, 3.20, 4.00, and 5.00 meters on the 5m setup. The time difference between the initial LED flash and the appropriate pushbutton press was recorded as the reaction time of that trial. This information was collected through the serial monitor on the Arduino IDE and converted into a comma separated values file for analysis. Incorrect button presses did not count as a trial. For each invalid trial resulting from an incorrect button press, an additional trial was added for a complete data set of 200 valid trials. In regards to the lattice data, the flashes were spaced throughout depths of 0.6m, 1.2m, 1.8m, and 2.4m and eccentricities of ±40°, ±20°, and 0°.

## Acknowledgments

A total ~60 students helped us as participants in various stages of RT experiments in 2020 – 2022. We thank them for their kind commitment.

This work was in part supported by the Dean’s office of life science, Dean’s office of physical science, Chair’s office of the department of physics and astronomy, and the Instructional Improvement grant by the Center for the Advancement of Teaching, all at the University of California, Los Angeles.

## Contributions

KA developed the concept and supervised the project. IB, AK, DS, MB assisted in coordinating researchers and participants under the supervision of KA. NY, IB, AK, TM, JC developed the Arduino protocol. NY developed the initial experiment. AK, IB, NY, TM, UA, JC, PC, MD, KT, SN constructed the hardware at UCLA Knudsen Hall. Many other undergraduate students constructed and operated hardware individually for data taking. TM, UA, BT wrote the MatLab analysis code. DS, IB and BT analyzed the data. DE and EM managed the IRB. IB and MB oversaw the writing of the original manuscript. KA and AB edited and proofread.

